# Divergence in neurodivergent brains: mapping the relationship between developmental trajectories in functional connectivity and clinical outcomes

**DOI:** 10.64898/2026.09.23.753857

**Authors:** Sara Saljoughi, Kathleen Lyons, Taylor Heffer, Mehran Ebrahimi, Ryan Stevenson, Bobby Stojanoski

## Abstract

Brain network development differs between neurodivergent (ND) and neurotypical (NT) youth, but its developmental trajectory, and relationship to clinical outcomes, remains unclear. We examined age-related changes in functional network connectivity and clinical symptoms in 861 participants (5–21 years) from the Healthy Brain Network, including NT youth and those with Neurodevelopmental Disorders (NDDs) and generalized anxiety disorder (GAD). Functional connectivity during movie-watching was modeled using generalized additive models across eight large-scale networks to characterize nonlinear developmental trajectories. Divergence in connectivity profiles between ND and NT groups first emerged during late childhood and were largest during adolescence. Importantly, divergence in connectivity patterns followed a similar trajectory path, but preceded, divergence in clinical outcomes. These findings identify adolescence as a sensitive period when differences in patterns of brain activity and clinical differences become increasingly pronounced, highlighting a potential window for targeted intervention.

## 1. Introduction

The brain undergoes massive reorganization during childhood and adolescence, including the emergence and segregation into increasingly dissociable cortical functional networks ^1^. Neural reconfiguration from childhood to adolescence coincides with significant improvements to various social-cognitive abilities that are fundamental for navigating complex and often changing social environments ^2,3^. Although large-scale reorganization mostly stabilizes during adolescence, continued refinement across distributed networks during this period ^4^ is likely not linear ^2^, and variations in these developmental trajectories may contribute to the clinical symptomatically profiles observed in neurodivergent youth ^5^.

Neurodivergent (ND) youth, including those with Autism Spectrum Disorder (ASD), Attention-Deficit/Hyperactivity Disorder (ADHD), and co-occurring Autism and ADHD (AuDHD) and Generalized Anxiety Disorder (GAD), face developmental challenges that their neurotypical peers typically do not, which can affect well-being across the lifespan ^6^. These include circumscribed challenges in executive functioning and learning, to more pervasive disruptions in social competence ^7,8^. There is also evidence that neurodivergent youth appear to have distinct cortical organization across different structural and functional networks relative to neurotypical children (NT) in childhood and adolescence. For instance, altered subcortical, sensorimotor, and frontal circuits are found in Autistic children ^9^; frontal dysconnectivity is linked to impairments in attention and reward processing in children with ADHD ^10^, and disruptions in fronto-amygdala and fronto-limbic connectivity are reported in children with anxiety disorders ^11^. It is plausible that symptomology and socio-cognitive challenges experienced by neurodivergent youth groups could be associated with deviations in the developmental trajectory of cortical network organization ^12^.

Despite advances in characterizing brain development in the neurotypical population, trajectories in neurodivergent youth and their relation to symptom scores remain poorly defined, with inconsistent findings particularly in brain regions involved in socio-cognitive processing ^13–16^. For instance, some studies have reported autistic adolescents show early-life within default mode network (DMN) hyperconnectivity that resolves in adulthood ^17,18^. In contrast, others have identified childhood hyperconnectivity between the DMN, frontoparietal, and salience networks in Autistic youth, which either persisted into adulthood or decreased to hypoconnectivity in adulthood ^19^. These inconsistencies likely arise from large differences in statistical and methodological approaches, such as inconsistent age binning, or relying on parametric (i.e., linear) models or restricted nonlinear approaches to characterize brain networks patterns, despite the inherently nonlinear and in flux nature of brain development ^20^. Moreover, variability in the ecological validity ^21^ of experimental paradigms may be contributing to inconsistent findings. For instance, traditional task-based paradigms often rely on simplified, artificial stimuli that may fail to capture the complexity of real-world environments ^22^, whereas task-free paradigms, such as resting-state fMRI, while valuable, do not fully capture the dynamic nature of social-cognitive processing of real-world information ^23^. Naturalistic paradigms and a more flexible modeling framework are needed to capture subtle changes in brain network development and how they might be related to clinical measures.

The current study addressed these gaps by applying Generalized Additive Models (GAMs) to characterize the developmental trajectories of key brain networks in a large cross-sectional sample of neurodivergent (ASD, ADHD, AuDHD, GAD), and NT youth during a naturalistic movie-watching paradigm. Movie-watching captures the complexity of real-world experiences by providing continuous, dynamic, and socially rich stimuli that naturally engage social-cognitive processes that resting-state and non-narrative paradigms do not ^22^. Additionally, GAMs provide a data-driven framework for modeling both linear and nonlinear age-related changes in functional connectivity^24^, while identifying developmental inflection points at which participant groups begin to diverge. Integrating clinical measures within these developmental trajectories helps characterize divergent patterns of brain network maturation across neurodivergent and neurotypical youth and identify potential developmental windows of heightened vulnerability associated with neurodivergent profiles. Specifically, this study aimed to 1) characterize differences in functional brain network development between neurodivergent groups, 2) identify the brain networks in which these differences are most pronounced, 3) demarcate when differences in key inflection points emerge and how long they persist and 4) examine whether divergence patterns in brain connectivity mirror divergence in clinical symptoms. Identifying deviations in the time course of cortical network developmental development across childhood and adolescence may help delineate sensitive developmental periods that can inform earlier diagnosis, more targeted interventions, and educational and social policies that better support neurodivergent individuals.

## 2. Methods and Materials

### 2.1. Participants

We used neuroimaging data made publicly available by the Healthy Brain Network biobank (releases 1–8) ^25^, an initiative of the Child Mind Institute. A total of 861 participants (327 females; *M* = 11.28 years, *SD* = 3.61; age range = 5–21 years) were included in this study. Inclusion criteria required a clinical diagnosis criterion of ASD, ADHD, comorbid ASD and ADHD, and GAD along with a NT (i.e., no diagnosis), and availability of movie-based task data. Diagnoses were determined using the HBN Diagnosis_ClinicianConsensus instrument, which allows for up to ten clinician-assigned diagnoses per participant. Participants were categorized into three diagnostic groups: NT, GAD, and neurodevelopmental disorders (NDD). The NDD group consisted of individuals with ASD, ADHD, and AuDHD (Table1). Group membership was based on at least one diagnosis matching the diagnostic category; however, participants were not excluded for the presence of co-occurring diagnoses; there are likely no “pure” clinical categories and, thus, this would be more representative of each clinical subpopulation. For example, a participant with at least one diagnosis of anxiety disorder would be in the GAD group despite potentially also having other non-NDD diagnoses. In the case where a participant was diagnosed with GAD *and* ADHD, ASD, or AuDHD, they would be included in the GAD group. Similarly, a participant with a diagnosis of a neurodevelopmental disorder would belong to the NDD group despite any co-occurring diagnosis other than GAD. For example, an Autistic participant diagnosed with any other condition except GAD would be in the NDD group. The large number of participants with a diagnosis of ADHD without a co-occurring condition allowed us to create an ADHD subgroup.

### 2.2. fMRI stimuli and processing

All imaging data were preprocessed through the Reproducible Brain Charts (RBC) project ^26^, an open resource supporting investigations of brain development and mental health. Structural MRI data were processed using FreeSurfer and sMRIPrep, while functional MRI data were preprocessed with the Configurable Pipeline for the Analysis of Connectomes (C-PAC)^27^, an open-source pipeline designed to ensure reproducible connectomics. To mitigate scanner- and site-related variability, imaging features were subsequently harmonized across datasets using CovBat-GAM ^28^. Age and sex assigned at birth were included as phenotypic variables, and participant demographics are detailed in Table 1. The Chesapeake Institutional Review Board approved the study, and details on the HBN biobank can be found here: http://fcon_1000.projects.nitrc.org/indi/cmi_healthy_brain_network/. The institutional review board at Ontario Tech University approved secondary analysis of the HBN data.

**Table 1.** Summary of participant demographics and diagnosis.

|  |  |  |
| --- | --- | --- |
| **Sex** | N(%males) | 861(62%) |
| **Age** | Age (mean (SD)), years | 11.28(3.61) |
| **Motion** | Mean framewise displacement (mean FD) | 0.13(0.078) |
| **Race%** | White/Caucasian | 52.26% |
|  | Black/African American | 13.43% |
|  | Hispanic | 10.13% |
|  | Asian | 2.81% |
|  | Others | 21.37% |
| **Diagnosis (N (%))** | NT | 227(26%) |
|  | GAD | 189 (21%) |
|  | NDD | 445(51%) |
|  | ASD | 143(16%) |
|  | ADHD | 414(48%) |
|  | AuDHD | 112(13%) |

Neuroimaging data was collected, included T1-weighted anatomical images, and functional MRI scans while participants watched a 10-minute clip from the animated film *Despicable Me*. This clip was chosen because it is engaging and provides a naturalistic context for investigating the neural mechanisms of brain development across diverse youth populations by encouraging viewers to follow character development, and interpret rich social interactions to understand the plot ^29,30^.

Functional connectivity matrices were generated for each participant using 58 seed regions spanning eight networks of interest: Visual (VIN), Somatomotor (SMN), Dorsal Attention (DAN), Salience (SAN), Limbic (LIN), Frontoparietal (FPN), Default Mode (DMN) from the Yeo 7-network parcellation (51 nodes) ^31^, and the Theory of Mind (ToM) network (7 nodes) ^32^. Functional connectivity was estimated by calculating Pearson correlations between the time series of each pair of regions of interest (ROIs). Correlation coefficients were standardized using z-scores and assembled into a 58 × 58 connectivity matrix for each participant. Mean connectivity values were then averaged within and between the eight networks, yielding an 8 × 8 matrix consisting of eight within-network correlations and 28 unique inter-network correlations.

### 2.3. Generalized additive models

We used GAMs to analyze age-related differences in both within and between-network connectivity across neurotypical and neurodivergence youth. GAMs provide a robust and flexible statistical framework for modeling complex, nonlinear relationships, as they do not impose predefined functional forms between predictors and outcomes, unlike traditional regression approaches ^33,34^. Alternatively, these relationships are derived from the data during model estimation, eliminating the need for manual refinement and variable selection as enhancing both efficiency and precision in large-scale connectivity analyses ^35,33^. To prevent overfitting, GAMs employ cross-validation and regularization techniques, allowing them to effectively capture nonlinear associations between age and functional connectivity.

We examined the relationship between age and functional connectivity within eight brain networks and across twenty-eight inter-network connections, separately for each group. To consider potential confounding influences, sex and mean framewise displacement (FD) ^36^ were included as covariates in the GAMs. Model significance within each group was evaluated for age-related effects, with p-values corrected for multiple comparisons using the Benjamini–Hochberg false discovery rate (FDR) procedure. In GAMs, EDF shows the complexity of the relationship between variables, for instance, an EDF around 1 indicates linearity, an EDF around 2 reflects a quadratic pattern, and an EDF around 3 represents a more complex, cubic-like relationship ^33^.

All GAMs were fitted using Restricted Maximum Likelihood (REML) estimation ^33,34^ as implemented in the **mgcv** package (version 1.8-42; Wood, 2017) within R (version 4.3.1; R Core Team, 2019). Visualization of model outputs and scatterplots was performed using the **ggplot2** package ^37^.

#### 2.3.1. Between groups contrasts: functional connectivity

To examine group-related differences in developmental trajectories of key brain networks, *post hoc* pairwise comparisons were performed on the predicted trajectories derived from the generalized additive models. These analyses were conducted using the **tidy-gam** package ^38^.

Significant between-group differences were identified as age ranges where the 95% confidence interval (CI) of the difference excluded zero. To account for potentially spurious results and increase the reliability of the findings, effects that spanned over 2.5 years were considered significant (significant periods are highlighted in gray (see Figure 2D; Supplementary Figure 1). The analysis was implemented at two levels: **Level 1** characterized developmental differences across the broad diagnostic groups, including NDD, GAD, and NT participants, and **level 2** subdivided the NDD group into three subgroups, ADHD, ASD, and AuDHD, each of which was then individually compared with the NT group to account for the heterogeneity encompassed within the NDD category.

Following the *post hoc* pairwise comparisons, all significant age windows for each group comparison were summed and consolidated for within and between connectivity profiles for each network of interest. Summary plots display significant between-group differences at specific age ranges within and between brain network, providing a comprehensive overview of developmental divergence across conditions at both levels. That is, the temporal extent of developmental differences (i.e., significant age windows) was represented as duration bars for each comparison, with the overall aggregated pattern shown in black to highlight the duration and overlap of effects across groups. To characterize the most prominent brain networks contributing to these differences, radar plots were generated for each comparison category as well as for the aggregated results within each analysis level. This framework helped identify the emergence of age-specific divergence in connectivity patterns across diagnostic groups. These developmental windows may reflect key inflection points at which differences between neurotypical and neurodivergent brain development become apparent, potentially pinpointing windows of vulnerability for divergent developmental trajectories.

#### 2.3.2. Between groups contrasts: clinical profiles

To compliment the functional connectivity analysis, we also explored differences in clinical measurements between neurotypical and neurodivergent individuals. Clinical presentation was based on three domains: ADHD-symptoms, measured using the Inattention and Hyperactivity/Impulsivity subscales of the Conners 3 Self-Report (C3SR)^39^; autism-related traits were measured using the Autism Spectrum Screening Questionnaire (ASSQ)^40^; and anxiety-related symptoms were measured using the parent report version of the Screen for Child Anxiety Related Emotional Disorders (SCARED)^41^. Scores were first z-transformed and absolute mean differences between neurodivergent and neurotypical groups were calculated for each clinical scale and aggregated across development using the following age bins: 5– 7, 7–9, 9–11, 11–13, 13–15, 15– 17, 17–19, and 19–23 years, (in age bins) to characterize overall patterns of group divergence.

## 3. Results

### 3.1. Diagnostic specific developmental trajectories of network connectivity

Within-group analyses revealed divergent patterns of functional brain network development. Of the 36 brain functional connectivity measures analyzed for each broad group (NT, NDD, and GAD), 9 showed significant age-related changes in the NT group, whereas 17 connections exhibited significant age-related changes in the NDD group, and 3 connections were significant in the GAD group (FDR-corrected α < 0.05) (Fig. 1). The curvature of the connectivity–age trajectories was characterized using the effective degrees of freedom (EDF) derived from the GAM models. In general, the NDD group exhibited a more prominent decreasing pattern of connectivity in several networks including FPN and ToM compared to the NT and GAD groups. For example, ToM–FPN showed a linear decreasing trend in the NDD group (EDF = 1, F = 19.06, *p* < 0.001), FPN–SMN showed a decreasing quadratic pattern in the NDD group (EDF = 2.02, F = 8.80, *p* < 0.001), and DAN–VIN showed a more complex nonlinear, cubic-like increasing pattern in the NT group (EDF = 3.57, F = 4.81, *p* = 0.008). Curvature differences, along with overall increasing or decreasing trends within and between all networks of interest are illustrated in Fig. 1.

**Figure 1:**
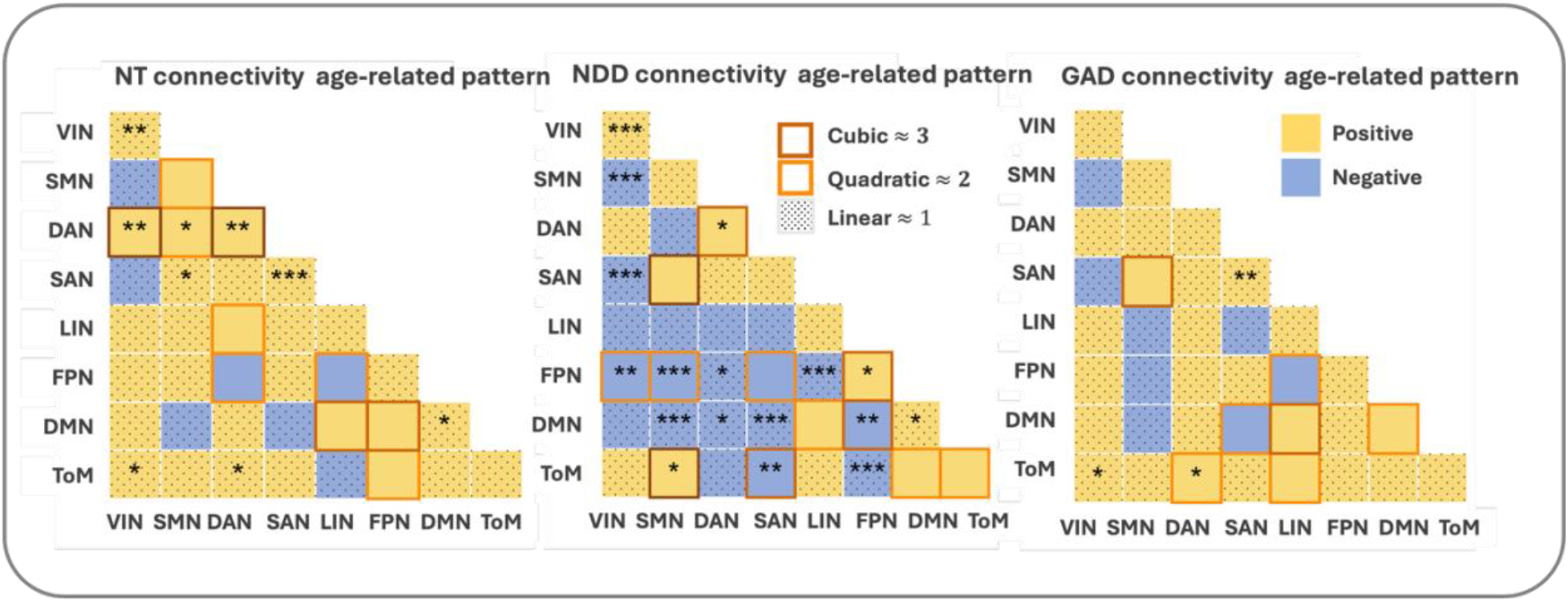
Age-related trajectories of functional brain networks. Curvature differences are displayed by brown, orange, and dotted markers for cubic, quadratic, and linear trajectories, respectively. Increasing and decreasing patterns are shown in yellow and blue. * Significant connections (* 0.05, ** 0.01, *** <0.001; FDR-corrected)

### 3.2. Divergent developmental trajectories of functional connectivity between diagnostic groups

Pairwise contrasts of the GAMs between the NDD, NT and GAD groups revealed connectivity profiles follow distinct developmental trajectories across diagnostic groups within the networks of interest.

NDD vs NT: This comparison yielded the most substantial differences primarily between the ages of 15 and 21 (Fig. 2A). During this period, we found those in the NT group exhibited stronger connectivity across several networks of interest, but most predominantly involving the FPN, including connectivity between FPN and DMN (NT > NDD, ages 14.9–20.2: Δ = 0.019–0.052; 95% CI > 0), FPM and ToM (NT > NDD, ages 14.9–21.5: Δ = 0.017–0.072; 95% CI > 0), FPN and SAN (NT > NDD, ages 16.2–19.5: Δ = 0.020–0.033; 95% CI > 0), and FPN and SMN (NT > NDD, ages 16.2–21.5: Δ = 0.036–0.088; 95% CI > 0). During the developmental window between 15 and 21 years of age, we also found the NT group showed stronger connectivity between ToM and VIN (ages 16.2–20.8: Δ = 0.029–0.054; 95% CI > 0), ToM and SAN (ages 14.2–20.2: Δ = 0.020–0.049; 95% CI > 0), and within SAN (ages 14.9–21.5: Δ = 0.016–0.049; 95% CI > 0). The NT group also showed stronger connectivity between DAN and SMN during the same developmental period, but also DAN and ToM (Δ = 0.017–0.066; 95% CI > 0), and DAN and DMN (Δ = 0.015–0.062; 95% CI > 0) that emerged earlier around 11 years old but extended until 21. Divergence in connectivity profiles emerged at earlier developmental stages, specifically between the ages of 5 and 11, whereby the NT group exhibited stronger connectivity within VIN (Δ = 0.01–0.004; 95% CI > 0) and LIN (ages 7.6–12.9: Δ = 0.02–0.018; 95% CI > 0).

**Figure 2:**
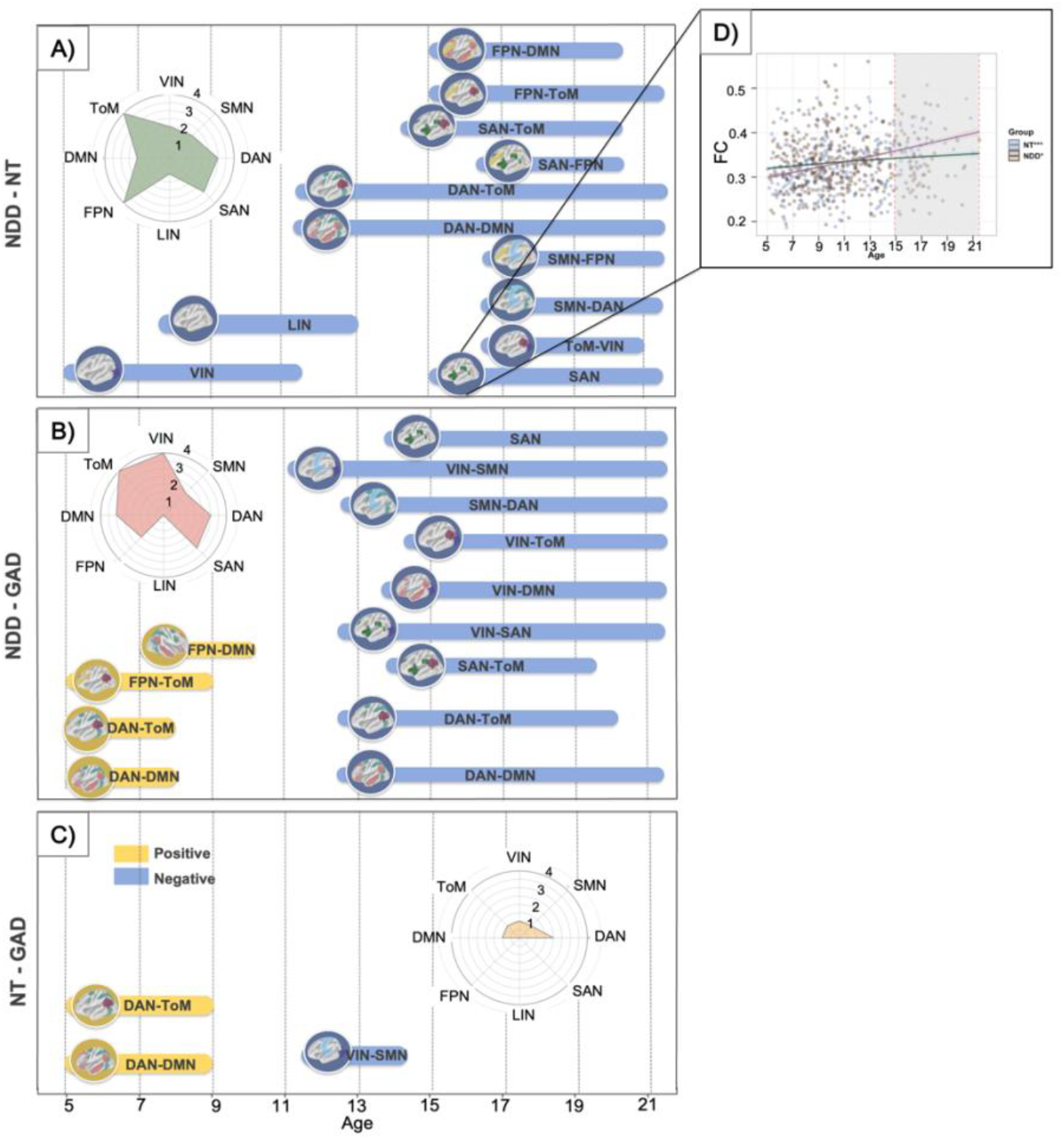
Overview of developmental differences in functional brain connectivity across diagnostic groups. (A–C) Pairwise comparisons of age-related functional connectivity trajectories between NDD and NT (A), NDD and GAD (B), and NT and GAD (C). (D) Example comparison showing connectivity differences between NT and NDD within the salience network and the corresponding statistically significant age intervals, highlighted in gray. Within each panel (A-C) spatial mapping of the total number of differences for each pairwise comparisons across networks.

NDD vs GAD: We also found several differences in the developmental trajectory of network connectivity between the NDD and GAD groups. These differences appeared at two distinct developmental periods. During the first period, between the ages of 5 to 10, we found the NDD group had stronger connectivity in between FPN and DMN (ages 7.2–9.8: Δ = 0.02–0.016; 95% CI > 0), FPN and ToM (ages 5.2–9.1: Δ = 0.03–0.017; 95% CI > 0), DAN and ToM (ages 5.2–7.9: Δ = 0.07–0.03; 95% CI > 0), and DAN and DMN (ages 5.2–7.9: Δ = 0.05–0.02; 95% CI > 0). In the second period, emerging at age 11 and extending to after 19 years of age, we found that the GAD group had stronger connectivity primarily in the VIN, specifically connections between VIN and SMN (ages 11.1–21.5: Δ = 0.034–0.1; 95% CI < 0), ToM (ages 14.3–21.5: Δ = 0.025–0.082; 95% CI < 0), DMN (ages 13.7–21.5: Δ = 0.02–0.085; 95% CI < 0) and SAN (ages 12.4–21.5: Δ = 0.023–0.064; 95% CI < 0). We also found stronger connectivity between DAN and DMN (GAD > NDD, ages 13.04–21.5: Δ = 0.02–0.09; 95% CI < 0), ToM (GAD > NDD, ages 13.04–20.2: Δ = 0.024–0.056; 95% CI < 0), and SMN (GAD > NDD, ages 12.4–21.5: Δ = 0.03–0.1; 95% CI < 0). See supplementary material for complete set of results.

NT vs GAD: This comparison yielded the fewest differences. We observed stronger connectivity between DAN and ToM (Δ = 0.07–0.03; 95% CI > 0) and DAN and DMN (Δ = 0.04–0.02; 95% CI > 0) for the NT group between the ages of 5 and 9. Later in development, between the ages of about 11 and 14, we found connectivity between VIN and SMN was stronger in the GAD group relative to the NT group (Δ = 0.037–0.041; 95% CI < 0).

Our results suggest network maturity follows a similar trajectory from childhood to adolescence for those in the NT and GAD groups, whereas youth with NDDs exhibit increasingly divergent connectivity patterns from youth in the GAD and NT groups during the same period. A summary of network connectivity that diverged most prominently across development are represented as radar plots (Fig. 2A–C). For contrasts between NDD and NT, we found the most frequent occurring differences included connections within and between FPN and ToM (4 instances each) (Fig. 2A). Across all networks comparing NDD and GAD, the most frequent occurring differences included connectivity within and between the VIN and ToM networks (4 instances each) (Fig. 2B), whereas divergence between NT and GAD were restricted to the DAN network (2 instances) (Fig. 2C). Complete set of the GAMs results isolating age-related differences for all within and between networks across each group-wise comparisons are provided in the Supplementary Material.

### 3.3. Transdiagnostic divergence of functional connectivity

To better understand the timing and duration of these differences we aggregated all significant time points (i.e., age ranges) emerging from the GAMs across all within and between networks between the diagnostic groups (NT, NDD, and GAD). The result is a global timeline of emerging brain-wide functional connectivity differences across all groups. We found a small initial peak in early to late childhood (approximately ages 5–9), followed by a sharp increase beginning around age 12 and peaking between approximately 16 and 20 (Fig. 3A). Notably, as most of these differences were driven by the NDD group, these patterns may indicate adolescence as a sensitive developmental period associated with the onset of cortical network maturation divergence, which is particularly prominent with NDDs. During these global periods of divergence, across all group level comparisons, the strongest effects were found in the ToM network, recurring across 9 connections, followed by DAN, VIN, SAN, and FPN (exhibited across 8, 7, 6, and 6 networks, respectively), highlighting divergent developmental trajectories within the social brain (Fig. 3B).

**Figure 3:**
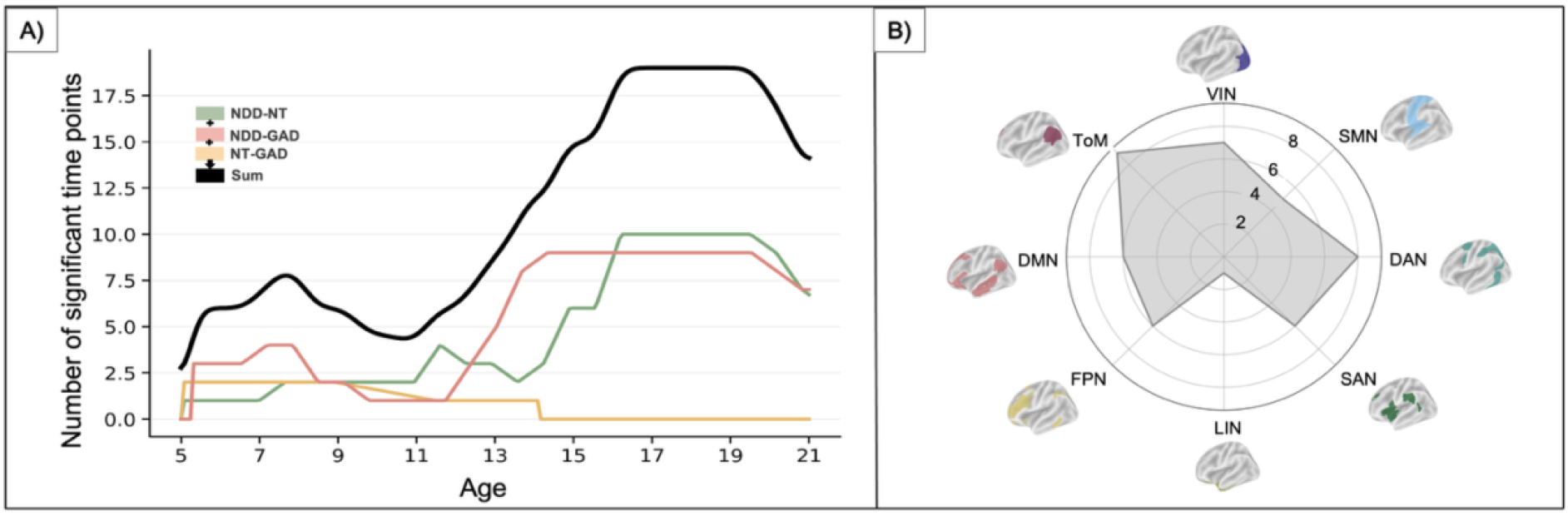
(A) Frequency of significant age points for each pairwise comparison (green: NDD–NT; red: NDD–GAD; yellow: NT–GAD) and their overall sum (black). (B) Spatial mapping of the total number of differences across all broad diagnostic pairwise comparisons across networks.

### 3.4. NDD subgroup analysis: ASD, ADHD, and AuDHD

Each NDD subgroup was separately compared with the NT group, following the same analytical framework outlined above. The subgroup analysis revealed significant differences between ADHD and NT (12 networks), ASD and NT (11 networks), and AuDHD and NT (7 networks), as outlined in Fig. 4A. Differences were most prominent in the ToM network (with additional effects in DMN, FPN, DAN, and VIN) for youth with ADHD; in DAN (along with VIN, SAN, and FPN) for autistic youth; and in VIN and FPN for youth with AuDHD. We observed a similar pattern of increasing connectivity differences beginning around age 7 and extending to approximately 11 (late childhood to early adolescence), followed by a second, larger window between ages 16 and 21 (mid- to late adolescence) (Fig. 4B). For the complete set of results see the Supplementary Material.

**Figure 4:**
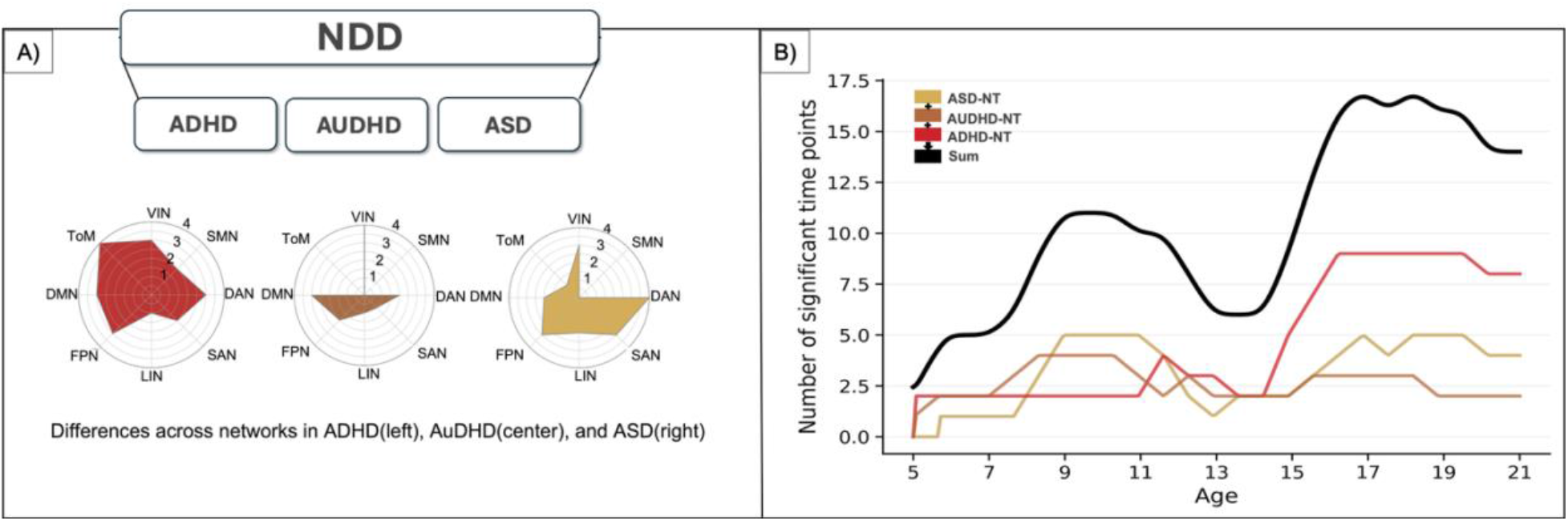
(A) Stage 2 NDD subgroup analysis. differences across networks for comparisons between ADHD, ASD, and AuDHD with NT. (B) Aggregation of significant age points across all NDD subgroup analyses.

### 3.5. Divergence in clinical profiles across development

We examined differences between neurodivergent and neurotypical individuals across three clinical scales. We found clinical scores across the three groups first diverged around age 9 with differences persisting until about age 15. We also found a later and larger period of divergence between the ages of 17 and 21 (mid- to late adolescence; Fig. 5). Notably, this temporal pattern aligns closely with the time windows of divergence in functional connectivity, however, with a short delay. This suggests divergence in connectivity profiles changes appear to precede differences in clinical presentation. Detailed age-bin results are reported in the Supplementary Material.

**Figure 5:**
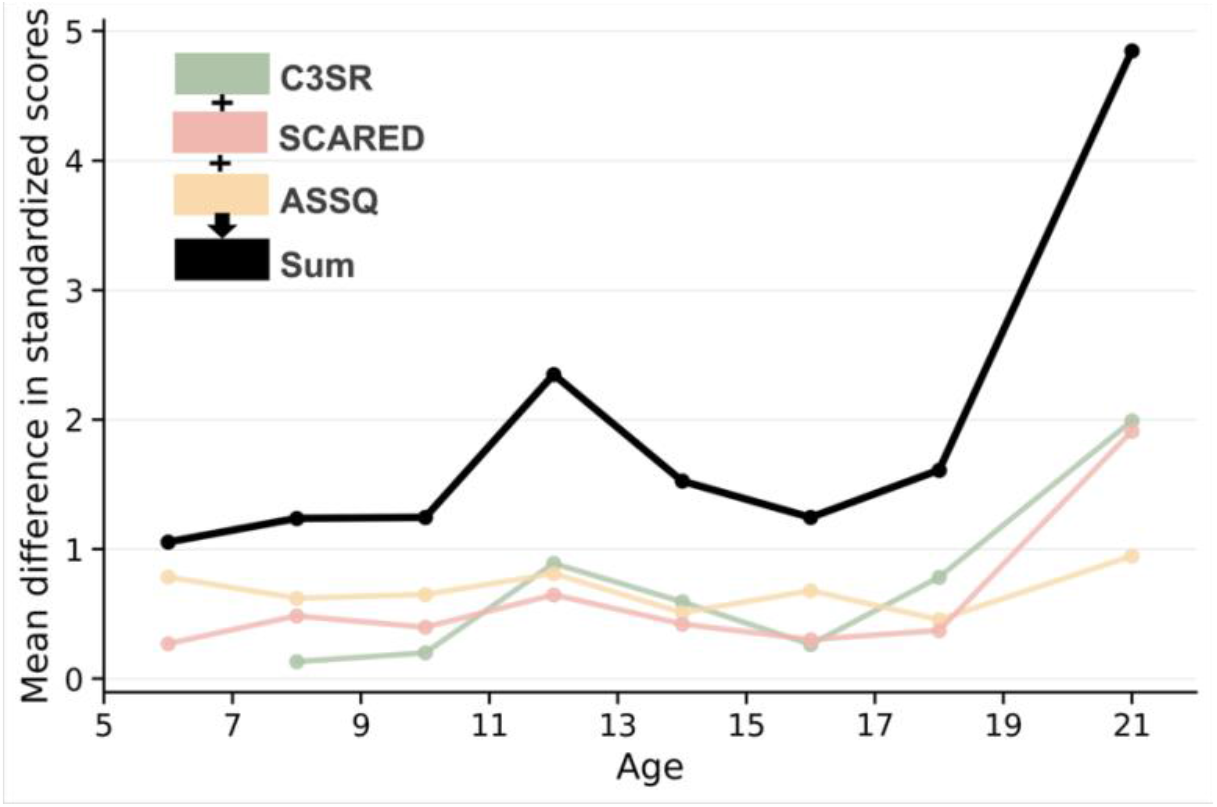
Age-related divergence in clinical profiles between neurotypical and neurodivergent participants. Group divergence was defined as the absolute difference in mean z-transformed scores between neurotypical and neurodivergent participants within each age bin.

## 4. Discussion

Through a transdiagnostic lens, we mapped sensitive developmental periods in both neural connectivity and clinical profiles between neurodevelopmental disorders (i.e., ASD, ADHD, and AuDHD), a developmental psychiatric condition (GAD), and typical neurodevelopment. We found that differences in functional connectivity between neurodivergent and neurotypical groups followed similar developmental trajectories, but directly preceded, divergences in psychiatric profiles. Specifically, cortical connectivity differences first emerged between early and late childhood, then widened during adolescence. These periods of maximal divergence were most pronounced in the Theory of Mind (ToM) network, followed by the dorsal attention (DAN), visual (VIN), salience (SAN), and frontoparietal (FPN) networks.

Interestingly, differences in the onset and duration of functional connectivity were largest between youth in the NDD group relative to those in the GAD and neurotypical groups with large differences in both within and between connectivity across the brain. In contrast, those in the NT vs. GAD groups differed relatively weakly only in connectivity between DMN with DAN and ToM, and between VIN and SMN. This suggests network maturity follows a similar trajectory from childhood to adolescence in the NT and GAD groups, while youth with NDDs exhibit increasingly divergent connectivity patterns relative to both groups over the same period. This finding is consistent with recent meta-analytic evidence suggesting that NDDs show an early peak age of onset around 5.5 years ^5,42,43–45^, whereas GAD shows a later peak age of onset around 15.5 years ^46^, which may reflect different developmental sensitive periods within different diagnoses.

A key factor in understanding vulnerability in neurodevelopment is identifying sensitive periods during which neurodivergent profiles show the greatest divergence from neurotypical trajectories. Here, we observed a small initial peak in divergence during early to late childhood, followed by a rapid increase beginning around age 12 and peaking between approximately 16 and 20 years across all networks and broad diagnostic groups. Intriguingly, this developmental pattern can be interpreted within the framework of the two-hit model, originally introduced to explain cancer development ^47^, and later extended to broader developmental and neurobiological processes ^6,48,49^. According to this model, the first hit, occurring in childhood, likely reflects early disruptions to neural development and genetic influences that fundamentally alter developing neural circuits. These compromised circuits, in turn, contribute to poorer behavioral outcomes and atypical neural organization later in development, rendering the brain more vulnerable to a second hit, which has been suggested to occur during adolescence ^6^. The two-hit model applied to autistic individuals, argues that the combined pressures of adolescent-specific developmental demands and pubertal hormonal changes, along with vulnerabilities in neural development arising from the first hit, cascade in a secondary hit that compromised neural circuitry may be unable to accommodate ^6^. Our findings align with the two-hit framework; the initial small increase observed during childhood may reflect the first hit, whereas the subsequent rapid increase during adolescence may correspond to the second hit. That is, the heightened divergence observed during adolescence may reflect the cumulative effects of earlier changes to neural connectivity profiles, hormonal changes, and environmental influences during the first hit. As a result, early emerging differences in neural development may become increasingly pronounced with age, limiting the acquisition of critical skills and behaviours needed to successfully navigate the transition from childhood to adolescence and, ultimately, to adulthood.

Examining specific NDDs, we observed a similar two-hit pattern in divergence in connectivity profiles relative to the neurotypical group. Connectivity differences increased gradually from approximately late childhood to early adolescence, followed by a second and larger window of divergence from approximately from mid- to late adolescence and early adulthood. This pattern was more pronounced for the ADHD and ASD group than the AuDHD group. The recurrence of this developmental pattern suggests that adolescence may represent a potential “sensitive” window during which neurodevelopmental differences become increasingly amplified, however, co-occurring neurodevelopmental condition may follow a distinct pattern of neural development.

Likewise, we observed two temporal windows of divergence pattern in clinical profiles (across the three scales) between neurodivergent and neurotypical participants. That is, SCARED and C3SR showed an initial, modest divergence between late childhood and early adolescents (ages 9 to 15), followed by stronger divergence during mid-to-late adolescence (ages 17 to 21). ASSQ showed a similar but less pronounced pattern, which may be partly explained by masking or compensatory processes in autism-related symptom presentation ^50^. Our results suggest a similar, but slightly delayed clinical two-hit pattern complimenting the observed network connectivity analysis.

Although we observed consistent temporal windows of divergence, these effects were not uniformly distributed across the brain. Aggregating across broad diagnostic groups, the strongest effects in the ToM and DAN networks, highlighting the important role of the social brain in divergent developmental trajectories. These findings are consistent with social-cognitive and social-communicative challenges core to ASD, and have also been recognized in ADHD ^43,51,52^. Challenges with social functioning, such as, fewer reciprocal friendships, difficulties with social interaction ^53^, and communication ^54,55^ have also been linked to AuDHD. Moreover, recent evidence suggests GAD is associated with alterations in social cognition and theory of mind, particularly in the context of worry about others and negative social information ^56^.

Similarly, divergence across developmental windows were more pronounced in some networks than others for neurodevelopmental disorders specifically. That is, the strongest and most consistent differences between the NDDs vs. NT groups were observed in VIN, DAN, FPN and DMN networks. The prominent involvement of the VIN aligns with prior research suggesting that neurotypical adolescents become increasingly sensitive to the global properties of visual stimuli, in this case the movie, whereas this shift in visual sensitivity may be developmentally reduced or even absent in autistic adolescents ^6,57^, indicating altered maturation of visual-processing systems in neurodevelopmental conditions.

A transdiagnostic approach, one that integrates neurodevelopmental conditions, psychiatric disorders, and normative development, is essential for establishing a comprehensive framework to better understand the various trajectories of brain development ^5^. For instance, co-occurring conditions, such as AuDHD, have historically been overlooked in clinical and research settings, which may have created barriers to appropriate identification of brain development in these individuals. The current framework provides new insight into the unfolding of divergent developmental pathways across neurodevelopmental, psychiatric, and normative trajectories. Particularly, our results have important implications for identifying when and where developmental deviations emerge across neurodivergent conditions. For instance, although adolescence is widely recognized as a period of heightened neural plasticity and developmental reorganization ^58^, our result extends this view by suggesting that this window may be especially consequential for neurodivergent youth. Identifying specific networks during this important window may help inform the timing of interventions, educational accommodations, and social supports, ensuring that neurodivergent individuals receive appropriate support during a period of rapid neural, cognitive, and social change.

## Acknowledgements

We would like to thank the Child and Mind Institute for designing and collecting data for the Healthy Brain Network Biobank. We would also like to thank the children, adolescents, and their families for taking the time to participate in studies conducted by the Child and Mind Institute. BS is funded by a Natural Sciences and Engineering Research Council of Canada Discovery grant (RGPIN-2020-05042), a CIHR Project Grant (487850), a Canadian Foundation for Innovation John R. Evans Leaders Fund (42163), and a SSHRC Insight Grant (435-2017-0936). RAS is funded through two NSERC Discovery Grants (RGPIN-2017-04656 & RGPIN-2024-06233), two SSHRC Insight Grants (435-2017-0936 & 435-2024-1375), a CIHR Project Grant (487850), the University of Western Ontario Faculty Development Research Fund, and a Canadian Foundation for Innovation John R. Evans Leaders Fund (37497), and through a grant from the Canada First Research Excellence Fund (OurBrainsCAN).

## Supplementary document

Detailed information regarding the pairwise diagnostic comparisons, including network-specific differences across groups, is provided in this supplementary document.

**Figure A.**
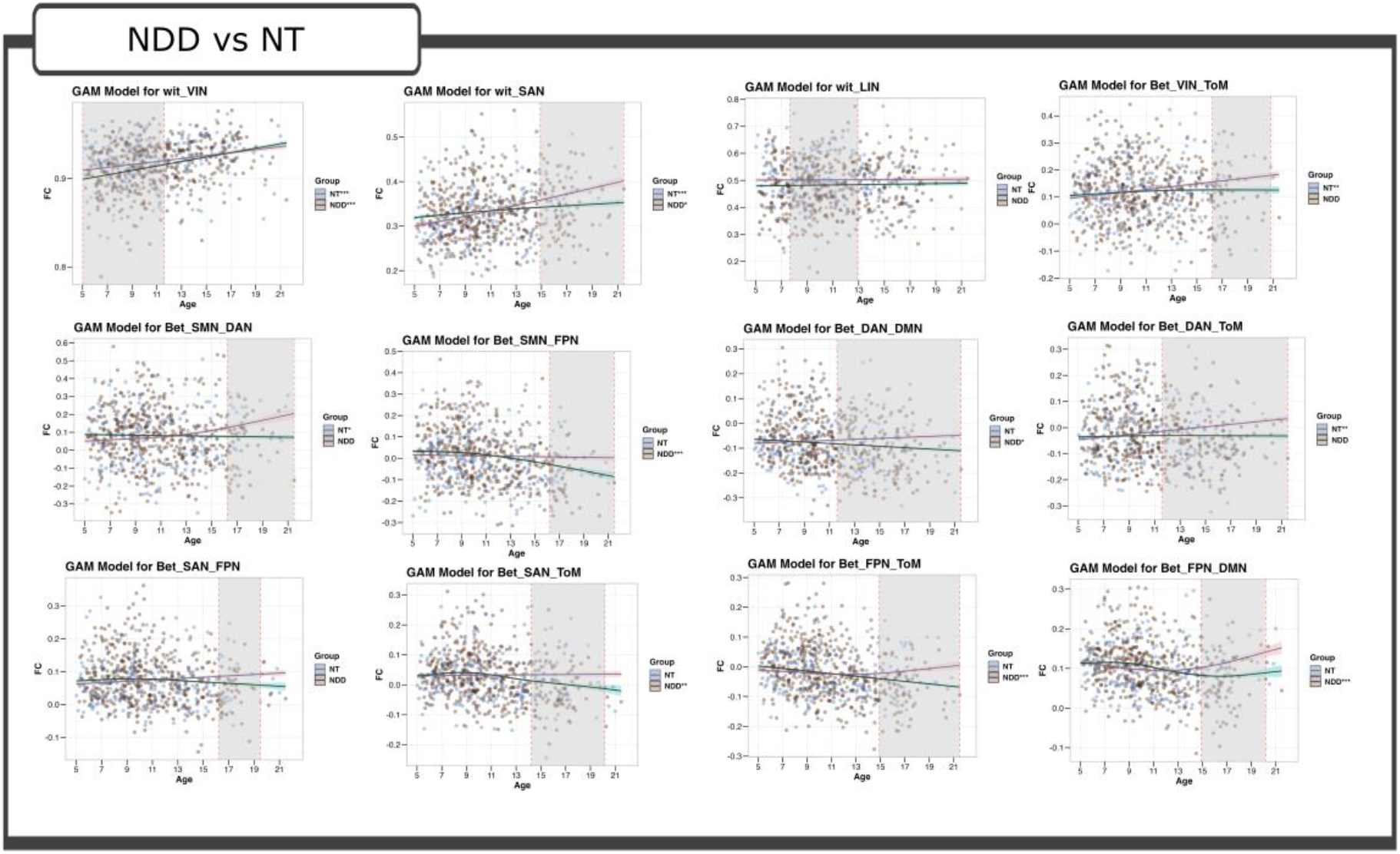
Detailed network-level results of the pairwise comparison between NT and NDD groups.

**Table A.** Detailed network-level results of the pairwise comparison between NT and NDD groups.

| Network | NT |  |  | NDD |  |  | Significant interval lower bound |  |  | Significant interval upper bound |  |  |
| --- | --- | --- | --- | --- | --- | --- | --- | --- | --- | --- | --- | --- |
|  | edf | F-value | p-value | edf | F-value | p-value | Delta | Age | 95%CI | Delta | Age | 95%CI |
| VIN | 1 | 13.22 | 0.0002 | 1 | 47.10 | <0.001 | 0.009 | 5.02 | [0.002, 0.017] | 0.004 | 11.6 | [0.0003, 0.008] |
| SAN | 1.16 | 24.54 | <0.001 | 1.28 | 3.38 | 0.03 | 0.016 | 14.9 | [0.001, 0.03] | 0.048 | 21.5 | [0.012, 0.08] |
| LIN | 1 | 0.01 | NS | 1 | 0.2 | NS | 0.02 | 7.6 | [0.0004, 0.04] | 0.0185 | 12.9 | [0.001, 0.035] |
| VIN-ToM | 1 | 8.65 | 0.003 | 1.4 | 0.6 | NS | 0.03 | 16.2 | [0.0002, 0.05] | 0.054 | 20.8 | [0.0004, 0.1] |
| SMN-DAN | 1.7 | 4 | 0.01 | 1 | 0.17 | NS | 0.05 | 16.2 | [0.004, 0.09] | 0.13 | 21.5 | [0.02, 0.24] |
| SMN-FPN | 1 | 0.17 | NS | 2 | 8.05 | <0.001 | 0.03 | 16.2 | [0.003, 0.068] | 0.08 | 21.5 | [0.01, 0.165] |
| DAN-DMN | 1 | 1.4 | NS | 1 | 4.6 | 0.03 | 0.015 | 11.6 | [0.0005, 0.03] | 0.06 | 21.5 | [0.017, 0.107] |
| DAN-ToM | 1 | 8.48 | 0.003 | 1.22 | 0.04 | NS | 0.02 | 11.6 | [0.001, 0.03] | 0.06 | 21.5 | [0.016, 0.11] |
| SAN-FPN | 1 | 3 | NS | 2 | 1.4 | NS | 0.02 | 16.2 | [0.0003, 0.03] | 0.03 | 19.5 | [0.0009, 0.06] |
| SAN-ToM | 1 | 0.04 | NS | 2.7 | 4.6 | 0.001 | 0.02 | 14.2 | [0.002, 0.03] | 0.48 | 20.2 | [0.002, 0.09] |
| FPN-DMN | 2.4 | 2.17 | NS | 2.9 | 5.3 | <0.001 | 0.02 | 14.9 | [0.002, 0.03] | 0.05 | 20.2 | [0.003, 0.1] |
| FPN-ToM | 1.9 | 0.6 | NS | 1 | 18.4 | <0.001 | 0.017 | 14.9 | [0.00008, 0.04] | 0.07 | 21.5 | [0.002, 0.12] |

**Figure B.**
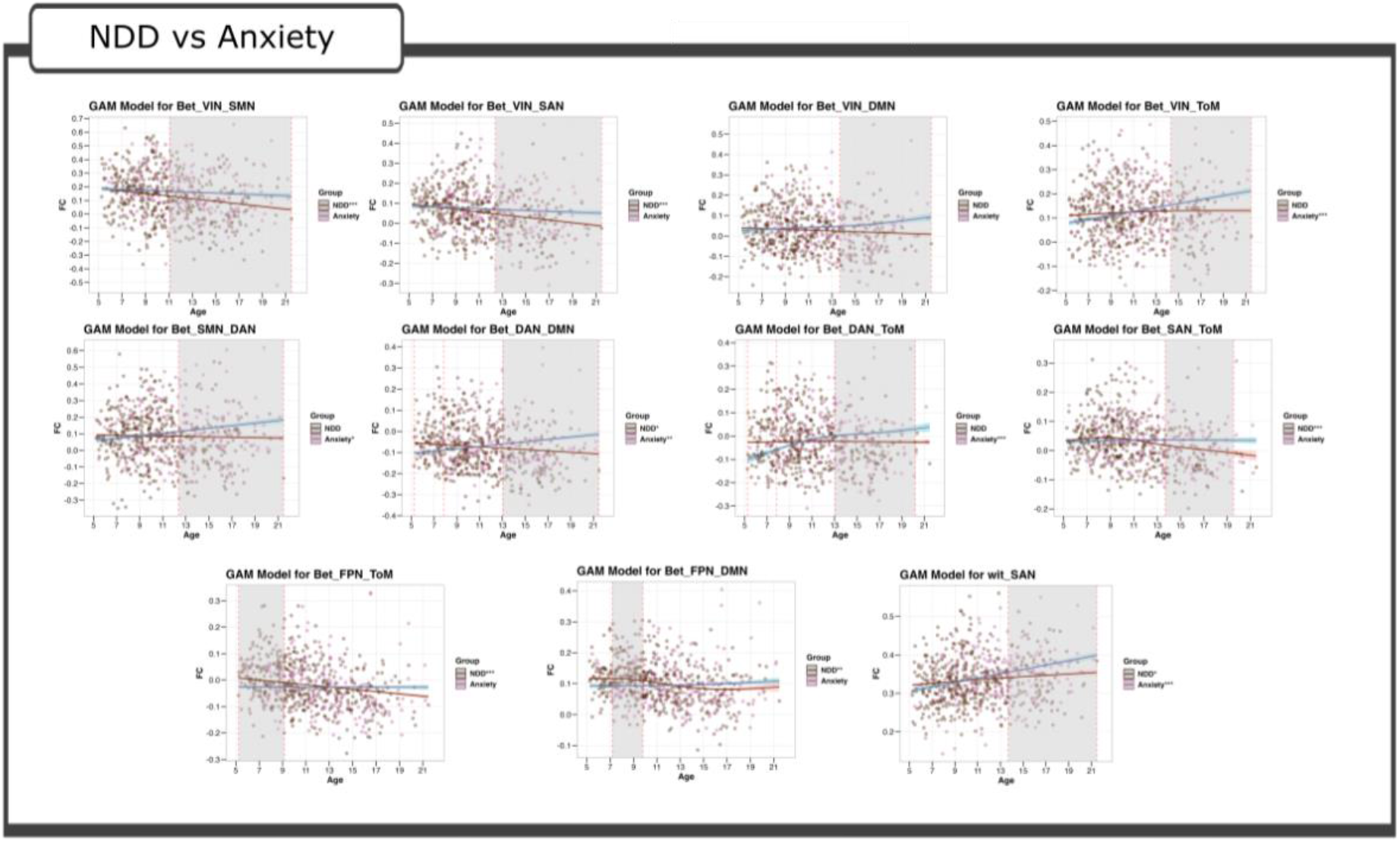
Detailed network-level results of the pairwise comparison between GAD and NDD groups.

**Table B.** Detailed network-level results of the pairwise comparison between GAD and NDD groups.

| Network | NDD |  |  | GAD |  |  | Significant interval lower bound |  |  | Significant interval upper bound |  |  |
| --- | --- | --- | --- | --- | --- | --- | --- | --- | --- | --- | --- | --- |
|  | edf | F-value | p-value | edf | F-value | p-value | Delta | Age | 95%CI | Delta | Age | 95%CI |
| SAN | 1.2 | 3.3 | 0.03 | 1 | 17.8 | <0.001 | 0.01 | 13.7 | [-0.02, -0.001] | 0.04 | 21.5 | [-0.07, -0.01] |
| VIN-SMN | 1 | 16.4 | <0.001 | 1 | 0.8 | NS | 0.03 | 11.1 | [-0.06, -0.003] | 0.1 | 21.5 | [-0.18, -0.018] |
| VIN-SAN | 1 | 17.9 | <0.001 | 1 | 0.8 | NS | 0.02 | 12.4 | [-0.04, -0.002] | 0.06 | 21.5 | [-0.11, -0.009] |
| VIN-DMN | 1 | 1.7 | NS | 1.6 | 1.3 | NS | 0.02 | 13.7 | [-0.04, -0.0006] | 0.08 | 21.5 | [-0.15, -0.02] |
| VIN-ToM | 1.4 | 0.6 | NS | 1 | 14 | <0.001 | 0.02 | 14.3 | [-0.04, -0.002] | 0.08 | 21.5 | [-0.14, -0.023] |
| SMN-DAN | 1 | 0.2 | NS | 1 | 5.7 | 0.01 | 0.03 | 12.4 | [-0.05, -0.001] | 0.1 | 21.5 | [-0.18, -0.034] |
| DAN-DMN | 1 | 6 | 0.01 | 1 | 8 | 0.005 | 0.04 | 5.25 | [0.013, 0.08] | 0.02 | 7.8 | [0.0002, 0.05] |
| DAN-DMN | 1 | 6 | 0.01 | 1 | 8 | 0.005 | 0.01 | 13 | [-0.03, -0.002] | 0.09 | 21.5 | [-0.13, -0.04] |
| DAN-ToM | 1.2 | 0.06 | NS | 2.2 | 6.3 | <0.001 | 0.07 | 5.25 | [0.16, 0.13] | 0.03 | 7.8 | [0.004, 0.06] |
| DAN-ToM | 1.2 | 0.06 | NS | 2.2 | 6.3 | <0.001 | 0.02 | 13 | [-0.04, -0.002] | 0.05 | 20.1 | [-0.11, -0.002] |
| SAN-ToM | 2.8 | 5 | <0.001 | 1.2 | 0.04 | NS | 0.018 | 13.7 | [-0.03, -0.0005] | 0.04 | 19.5 | [-0.08, -0.002] |
| FPN-DMN | 2.6 | 5.1 | 0.001 | 1.3 | 0.5 | NS | 0.02 | 7.2 | [0.002, 0.04] | 0.016 | 9.7 | [0.001, 0.03] |
| FPN-ToM | 1 | 17.7 | <0.001 | 1 | 0.01 | NS | 0.03 | 5.25 | [0.006, 0.06] | 0.017 | 9.14 | [0.0005, 0.03] |

**Figure C.**
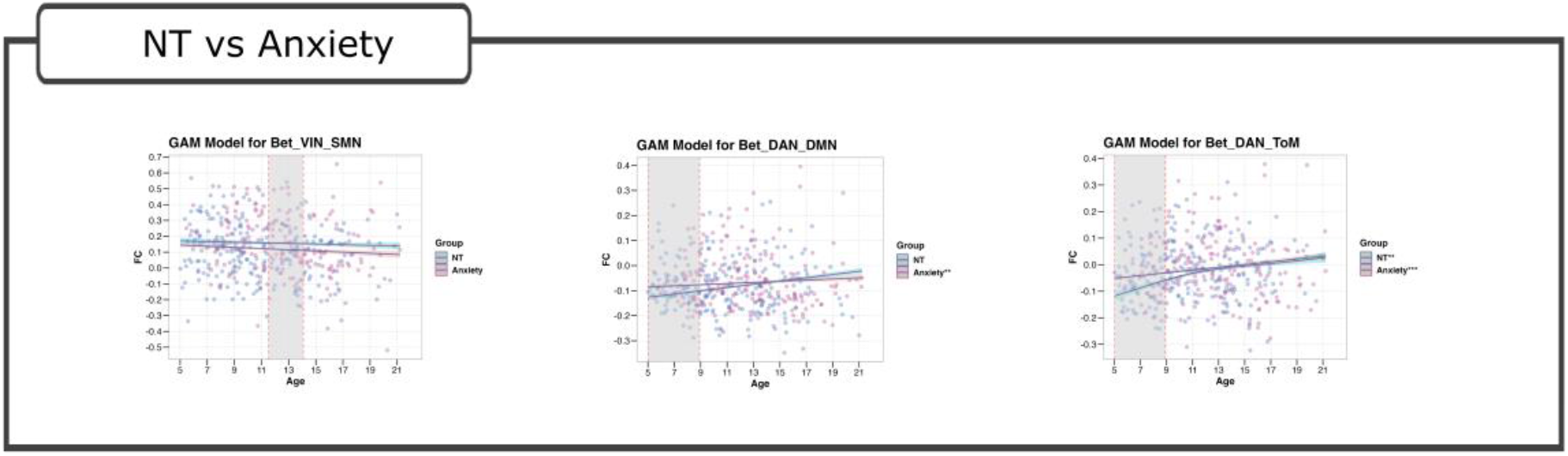
Detailed network-level results of the pairwise comparison between NT and GAD groups.

**Table C.** Detailed network-level results of the pairwise comparison between NT and GAD groups.

| Network | NDD |  |  | GAD |  |  | Significant interval lower bound |  |  | Significant interval upper bound |  |  |
| --- | --- | --- | --- | --- | --- | --- | --- | --- | --- | --- | --- | --- |
|  | edf | F-value | p-value | edf | F-value | p-value | Delta | Age | 95%CI | Delta | Age | 95%CI |
| VIN-SMN | 1 | 1.4 | NS | 1 | 0.2 | NS | 0.04 | 11.5 | [-0.072, -0.001] | 0.04 | 14.1 | [-0.08, -0.001] |
| DAN-DMN | 1 | 1.8 | NS | 1 | 10.3 | 0.001 | 0.04 | 5.02 | [0.001, 0.08] | 0.02 | 8.9 | [0.0001, 0.49] |
| DAN-ToM | 1 | 8.84 | 0.003 | 2 | 6.79 | <0.001 | 0.06 | 5.02 | [0.004, 0.12] | 0.27 | 8.9 | [0.0009, 0.05] |

**Table D.** Detailed age-bin results for mean (normalized) scores differences in clinical measures between neurodivergent and neurotypical participants.

| age_bin | C3SR |  |  | ASSQ |  |  | SCARED |  |  | Sum |
| --- | --- | --- | --- | --- | --- | --- | --- | --- | --- | --- |
|  | NT | ND | diff | NT | ND | diff | NT | ND | diff |  |
| (5, 7] | — | — | — | -0.48 | 0.30 | 0.784 | -0.416 | -0.146 | 0.270 | 1.054 |
| (7, 9] | 0.031 | 0.162 | 0.131 | -0.49 | 0.13 | 0.621 | -0.325 | 0.159 | 0.485 | 1.237 |
| (9, 11] | -0.109 | 0.089 | 0.198 | -0.573 | 0.075 | 0.649 | -0.341 | 0.054 | 0.396 | 1.245 |
| (11, 13] | -0.799 | 0.09 | 0.889 | -0.514 | 0.297 | 0.811 | -0.457 | 0.19 | 0.647 | 2.348 |
| (13, 15] | -0.4 | 0.193 | 0.593 | -0.423 | 0.085 | 0.509 | -0.18 | 0.241 | 0.422 | 1.525 |
| (15, 17] | -0.306 | -0.043 | 0.263 | -0.513 | 0.166 | 0.679 | -0.278 | 0.023 | 0.301 | 1.244 |
| (17, 19] | -0.552 | 0.23 | 0.783 | -0.237 | 0.218 | 0.455 | -0.313 | 0.058 | 0.371 | 1.610 |
| (19, 23] | -1.481 | 0.509 | 1.990 | -0.481 | 0.465 | 0.946 | -1.15 | 0.759 | 1.910 | 4.847 |

